# Protein sequence profile prediction using ProtAlbert transformer

**DOI:** 10.1101/2021.09.23.461475

**Authors:** Armin Behjati, Fatemeh Zare-Mirakabad, Seyed Shahriar Arab, Abbas Nowzari-Dalini

## Abstract

Protein profiles have many applications in bioinformatics. To construct the profile from a protein sequence, the sequence is aligned with database. However, sometimes there are no similar sequences with the query. This paper proposes a method based on pre-trained ProtAlbert transformer to predict the profile for a single protein sequence without alignment. The performance of transformers on natural languages is impressive. Protein sequences can be viewed as a language; therefore, we can benefit from using these models. We analyze the attention heads in different layers of ProtAlbert to show that the transformer can capture five essential protein characteristics of the family from a single protein sequence. These assessments are performed on the CASP13 dataset to find representative heads for each of five protein characteristics. Then, these heads are investigated on one thermophilic and two mesophilic proteins as case studies. The results show the significant attention heads for protein family properties extracted from a single protein sequence. This analysis led us to propose an algorithm called PA_SPP for profile prediction using only a single protein sequence as input. In our algorithm, we apply the masked language modeling method of ProtAlbert. The results display high similarity between the predicted profiles and HSSP profiles.

## 1. Introduction

Proteins consist of linear chains of twenty types of amino acids, each with different chemical properties. Proteins are the most versatile organic molecules in cells or living organisms and play critical roles in the body. The diversity of proteins functions is generally related to their diverse structures. The sequence of amino acids determines a unique protein tertiary structure which directly impacts its specific function^1^.

New sequencing technologies have led to an explosion in generating biological data such as protein sequences in the past two decades. UniProt^2^ Archive and Swiss-Prot^3^ databases contain most of the publicly available protein sequences globally. These sequences grow exponentially every few years^2^. Despite the strong interest in protein structure determination, there is currently a massive gap between the number of known sequences and experimentally determined structures deposited in the Protein Data Bank^4^ (PDB), highlighting the difficulties of structure elucidation^5^. Therefore, computationally predicting protein structure from the query sequence remains to be largely unsolved^6 7 8^. Homology modeling is a common approach for protein structure prediction. In this approach, homologous proteins of the query sequence are found by sequence comparison in a database. Then, a sequence profile is created to show the conservative and non-conservative regions in the homologous sequences ^9 10^.

Profiles are used in many bioinformatics problems. For example, they are applied to model protein families^11^, predict protein domains^12^, detect protein homology^13 14^, design proteins^15 16^, and identify orthologous genes and proteins^17^. Homology-derived Secondary Structure of Proteins (HSSP) database includes a sequence profile for each PDB protein. In HSSP, a Multiple Sequence Alignment (MSA) of putative homologs is prepared to construct a profile for each PDB protein. The list of homologous sequences is the result of an iterative database search in Swiss-Prot^18^. A well-defined profile can group information of similar sequences on conserved regions. It helps us to assign a query sequence to the family. This assignment is challenging when the query sequence length is short, or there is little similarity between this sequence and any sequences in the profile.

Protein structures are more conserved than protein sequences. Homologous proteins sharing a common evolutionary ancestor can have high sequence-level variations^19^, and when the protein sequence similarity is below 30% at the amino acid level, the alignment score usually falls into a twilight zone^20 21^. Therefore, simply comparing sequence similarities often fails to capture global structural and functional similarities of proteins.

Concerning the above discussion, improving the profile prediction methods to get more information about the sequence and families is an active research area in bioinformatics. In this paper, our primary goal is to predict a profile for query protein sequence without alignment using transformers.

In the following, we review the transformer-based models processing protein sequences. Proteins, as a linear chain of amino acids, can be viewed precisely as a language. Therefore, they can be modeled using Language Models (LMs) taken from Natural Language Processing (NLP). These LMs are used for biology identity representation and new prediction tools in various bioinformatics problems. The central concept behind this approach is to interpret protein sequences as sentences of characters (amino acids) and each character as a single word^22 23 24^. Recent research has shown that contextualized representations in NLP work well for contextual protein representation learning^25 26^. In the training phase, LMs learn to extract useful features from many samples and generate appropriate representations of these features ^27 28 29 30^. In these papers, architectures inspired by NLP are employed for protein processing. Also, pre-training tasks such as Masked-Language Modeling (MLM) and autoregressive generation are utilized to investigate protein-specific pre-training tasks.

One of the latest architectures that showed significant superiority over previous models is transformer^31^. Devlin et al.^32^ introduced a new language representation model based on transformers called Bidirectional Encoder Representations from Transformers (BERT). This model is designed to pre-train deep bidirectional representations from unlabeled text to create state-of-the-art models for a wide range of tasks. Bepler and Berge^33^ proposed a framework for mapping any protein sequence to a sequence of vector embeddings that encode structural information. Also, they defined a novel similarity measure between these arbitrary length vectors to learn useful position-specific embeddings. Similarly, Alley et al.^34^ used a Recurrent Neural Network (RNN) named UniRep to learn statistical representations of proteins and demonstrated that such representations predict the stability of natural and de novo designed proteins, as well as the quantitative function of molecularly diverse mutants. Rao et al.^35^ introduced TAPE as a new benchmark consisting of five relevant semi-supervised tasks for assessing such protein representation.

Elnaggar et al.^29^ trained two auto-regressive language models (Transformer-XL, XLNet) and two auto-encoder models (BERT, ALBERT) on data extracted from UniProt Reference Clusters (UniRef) datasets and Big Fat Database (BFD). They showed the effects of these pre-training models upon the success of the subsequent supervised training for predicting secondary structure, subcellular localization, and membrane-bound or water-soluble protein problems. Lu et al.^36^ applied the principle of mutual information maximization between local and global information as a self-supervised pre-training signal for protein embeddings to introduce a contrastive loss that trains an RNN to discriminate fragments from a source sequence versus randomly sampled fragments from other sequences. Min et al.^37^ introduced a novel pre-training scheme for protein sequence modeling called PLUS consisting of masked language modeling and a complementary protein-specific pre-training task, namely same-family prediction. They showed the advances of the PLUS on six out of seven protein biology tasks. Sturmfels et al.^38^ introduced a new pre-training task for protein sequence models. They used profile-hidden Markov models derived from MSAs as labels during pre-training for profile prediction. They utilized the model on a set of five downstream tasks for protein modeling and demonstrated that the model outperforms masked language modeling alone on all five tasks.

Although most previous studies on using transformer models for embedding protein sequences in different bioinformatics problems show acceptable results, they applied the model as a black box. Rogers et al.^39^ proposed an approach to interpret the heads and layers of transformers in NLP. Later, attention heads of some pre-trained transformers on proteins were interpreted to determine some of the characteristics of the proteins based on attention weights^40^. A probability was defined based on attention values to analyze each head in each layer to determine some protein characteristics.

Here, we further analyze more protein characteristics by proposing five new approaches to interpret attention weights in each head of different layers of ProtAlbert as one of the best transformers available. This assessment shows that some essential protein family features are captured in the attention heads of ProtAlbert given only a single protein sequence as input. The results of our analysis lead us to propose an algorithm for protein sequence profile prediction when only a single protein is given as input. We select pre-trained ProtAlbert because its efficiency enables us to run the model on longer sequences with less computation power while having similar performance with the other pre-trained transformers. Then, we propose five algorithms called RLH_NNI^i^, RH_SAA^ii^, RH_BBP^iii^, RH_PSS^iv^, and RH_PTS^v^ to analyze five protein characteristics, nearest-neighbor interactions, type of amino acids, biochemical and biophysical properties of amino acids, protein secondary structure, and protein tertiary structure at attention heads in the layers of ProtAlbert, respectively.

For this assessment, we make a dataset by extracting 55 proteins from CASP13 whose sequences, experimental tertiary structures, and HSSP profiles are available. In addition, we perform our analysis on one thermophilic and two mesophilic proteins as case studies. We show that representative heads on each protein characteristic are shared between the CASP13 proteins and case studies.

After executing each of five transformer head analyzer algorithms, we reach the following results:

- RLH_NNI algorithm detects representative heads in the layers of the ProtAlberl model for interaction between amino acids located at *k* (1 ≤ *k* ≤ 5) distances on the protein sequence.
- RH_SAA algorithm finds specific heads for aspartic acid, glutamic acid, proline, tryptophan, and histidine.
- RH_BBP algorithm announces representative heads for amino acids classified based on R-group.
- RH_PSS algorithm identifies some heads which contain significant attention weights from helix to helix, coil to coil, and sheet to sheet.
- RH_PTS algorithm finds a representative head for protein contact map, which is a simple tertiary structure representation.

Generally, these analyses show that the representative heads of the pre-trained ProtAlbert on protein sequences can detect protein family features. This result leads us to propose an algorithm called PA_SPP^vi^ for sequence profile prediction by pre-trained ProtAlbert on each single protein sequence using MLM. In other words, PA_SPP predicts profile from a single protein sequence without using MSA. We compare the predicted profiles to the HSSP profiles. The result shows the high similarity between the predicted and HSSP profiles. PA_SPP algorithm can help the researchers to predict a profile similar to the HSSP profile while there are no similar sequences to the query sequence in the database for making the HSSP profile.

## 2. Material and Method

This section first introduces the basic definitions needed to interpret ProtAlbert as a transformer model. Next, we propose five algorithms for assessing the layers and heads of ProtAlbert to identify some protein characteristics. Then, our approach is illustrated for the sequence profile prediction problem in more detail. In the end, we introduce the dataset used for evaluation.

### 2.1 Notation and Definition

The sequence of protein *P* with *n* length is represented by:

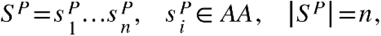

where *AA* = {*a*_1_,… *a*_20_} shows the set of amino acids. We define amino acid 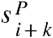 as a *k*-neighbor of 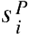 in sequence *S*^*P*^. The positive (negative) value of *k* shows that the position *i* attends from left to right (from right to left) of the sequence to find the neighboring amino acid at distance *k*.

In protein folding, the sidechain backbone of nearest-neighbor interactions may restrict the accessible conformations to a chain of protein^41^. Neighboring amino acids can be structurally categorized according to their separation in the primary sequence as proximal (1-4 positions apart) and otherwise distal^42^. For each protein *P*, *k*-neighbor interaction is defined based on interaction of each position *i* with position *i* + *k* on the sequence *S*^*P*^.

In addition to the effect of the nearest neighbor amino acids on protein folding, each amino acid has different biochemical and biophysical properties that can effectively determine the protein structure. Amino acids are classified based on R-group^vii^ into five classes 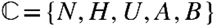 (see Table 1).

**Table 1:**
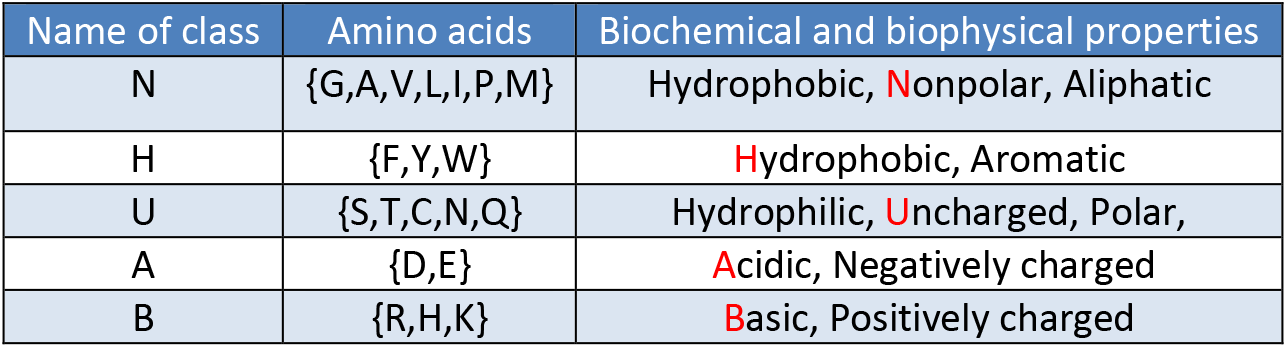
Classification of amino acids based on R-group: 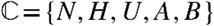.

The experimental structure of proteins can be extracted from PDB^viii^. Therefore, the 3D coordinate of each atom of amino acids in the protein 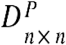 sequence is available. Here, we represent the tertiary structure of protein *P* with length *n*, by contact map as follows:

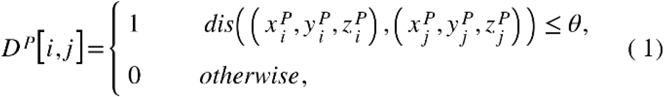

where *dis*(.,.) is the Euclidian distance and 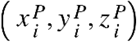 shows the 3D coordinate of the atom *c*_*α*_ for amino acid 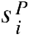 at position *i* of protein *P*. The value of *θ* is set 4.87 based on^43^. Each element *D*^*P*^[*i,j*] with value 1 indicates that two amino acids 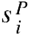 and 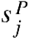 are in contact.

The secondary structure of protein is extracted from the tertiary structure using DSSP^ix^ software. This method provides eight classes, 3-helix, 4-helix, 5-helix, β-strand, β-bridge, turn, bend, and coil. Typically, the DSSP states are converted into three classes using the following convention. 3-helix, 4-helix, and 5-helix are considered helix (H). β-strand and β-bridge are displayed by a sheet (E). The rest of the states are shown as a coil (C). The secondary structure of protein *P* with length *n* is displayed as follows:

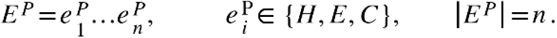

As mentioned in^44^, the secondary structure of each position in the protein sequence is dependent on its neighbors. The length of each type of regular secondary structure^45^ is about 6. We define the secondary structure matrix named 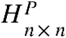 on protein *P* with length *n* as follows:

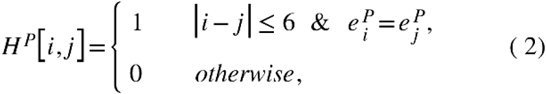

where *H*^*P*^[*i,j*] = 1 indicates the same secondary structure between two amino acids 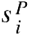 and 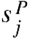 with distance less than 7 in sequence *S*^*P*^.

For each protein *P* with length *n* in PDB database, a profile named 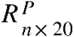 is extracted from the HSSP database^18^. In this database, there is an MSA of all available homologous sequences properly aligned to protein sequence *S*^*P*^. This MSA is constructed based on searching in the Swiss-Prot database considering the sequence family and structure. Each sequence of MSA is more than 30% identical to *S*^*P*^. Using MSA, the profile 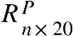 is generated where *R*^*P*^[*i,j*] shows the probability of amino acid *a*_*j*_ ∈ *AA* at position *i* of MSA.

In the following, we assume that dataset Δ = {*P*_1_,…,*P*_*t*_} includes *t* proteins where their sequences, experimental tertiary structures, and HSSP profiles are available.

### 2.2 ProtAlbert as a pre-trained transformer model on protein sequences

As described earlier, protein sequences can be viewed as a language, and therefore, we can benefit from using the models initially developed for natural languages. One of the latest architectures that showed significant superiority over previous models is transformers.

As it was mentioned, BERT^32^is a method of pre-training language representations. It means that after training a general-purpose language understanding model on a large corpus of text, the model can be used on downstream tasks. BERT is an example of auto encoding language modeling trained using MLM. During the training, 15% of the input is randomly masked, and the model is asked to predict the masked tokens. This process lets the model predicts the masked tokens based on the other available tokens. It shows that the model has a good idea about the language and the context. This self-supervised pre-training method, which means the labels are in the training corpus, got better results in many downstream tasks.

A year after BERT^32^, ALBERT^46^ was released by Google research that improved state-of-the-art performance in 12 NLP tasks. The main idea in the ALBERT was to allocate capacity more efficiently. They made two design changes to BERT, but the training process was MLM. First, while the input level embeddings need to be context-independent representations, the hidden-state embeddings need to take context into account. This was addressed by splitting the embedding matrix between a low dimension input-level embedding with length 128 and a higher dimension hidden-layer embedding with size 4096. The second critical change was removing redundancy and therefore increasing the capacity of the model to learn. Previously, it was observed that the various layers of BERT with different parameters in the model learned similar operations. This possible redundancy was eliminated in ALBERT by parameter sharing in different layers. These two design changes resulted in 90% parameter reduction compared to BERT with slightly decreased accuracy. However, this reduction allows scaling the hidden size from 768 in BERT to 4096 in ALBERT. It is shown that the bigger hidden layer embeddings can capture and represent the context better^46^.

We base our experiments on ProtAlbert, a transformer-based model on ALBERT architecture from the ProtTrans project^29^. ProtAlbert is pre-trained on 216 million protein sequences from the UniRef100 dataset. In this paper, we do not train or fine-tune the model. In the ProtAlbert model, the protein sequences are tokenized using a single space between each amino acid (indicating words), and each sequence is stored in a separate line (indicating sentences). Also, all non-generic or unresolved amino acids (B,O,U,Z) are mapped to the unknown token X. This model can process sequences with lengths of up to 40K, although this length is bound by the hardware capacity. The details of the ProtAlbert model are available in Table 2.

**Table 2:** ProtAlbert Parameters.

| Hyperparameter | ProtAlbert |
| --- | --- |
| Dataset | UniRef100 |
| Number of Layers | 12 |
| Hidden Layers Size | 4096 |
| Hidden Layers Intermediate Size | 16384 |
| Number of Heads | 64 |
| Positional Encoding Limits | 40K |
| Target Length | 512/2048 |

Our work contains two main parts, ProtAlbert analysis and profile prediction. For the first part, a protein sequence is given as an input to the ProtAlbert transformer. Then, we analyze and interpret the attention weights at attention heads in different layers to show that heads can capture some essential family features from a given single protein sequence. This assessment leads us to propose an approach for predicting a profile from a single protein sequence. In the second part, protein profile is predicted using ProtAlbert and masked token prediction. In other words, a protein sequence with some masked amino acids is fed to the model for predicting the most likely amino acids in the masked positions.

We choose ProtAlbert^29^ because its efficiency enables us to run the model on longer sequences with less computation power while having similar performance with ProtBert^29^, which is a great advantage. The ProtBert model is a pre-trained BERT-based language model with 420M parameters from the ProTrans project that has been trained on the same dataset as the ProtAlbert model with 224M parameters.

### 2.3 Proposed algorithms for analyzing ProtAlbert transformer to identify protein characteristics

In this sub-section, we propose five algorithms to analyze the attention heads and layers of ProtAlbert for finding the specific properties of proteins (see Table 3). This analysis is essential because it shows that ProtAlbert transformer can capture some biological features from a single protein sequence. It allows us not to apply the transformer as a black box but to select the ProtAlbert features specific to the bioinformatics problems.

**Table 3:** Five protein characteristics.

|  |
| --- |
| Nearest-neighbor interaction |
| Type of amino acids |
| biochemical and biophysical properties of amino acids |
| Protein secondary structure |
| Protein tertiary structure |

The input and output of this assessment are defined as follows:

- **Input**: Sequence 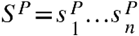 of protein *P*.
- **Output:** Extracting attention matrix 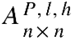 from ProtAlbert for each head *h* in layer *l* to interpret the properties of protein *P* displayed in Table 3.

ProtAlbert includes 12 encoder layers, and each encoder has 64 attention heads. Each protein sequence 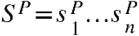 is given to the model as an input, then it goes through the encoder layers, and the attention mechanism in each layer generates output to go to the next layer.

For the input sequence *S*^*P*^ with length *n*, each attention head *h* (1 ≤ *h* ≤ 64) in the layer *l* (1 ≤ *l* ≤ 12) produces a matrix of positive attention weights named 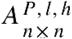. The value *A*^*P,l,h*^[*i,j*] shows the attention weight from amino acid 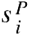 to 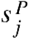 in layer *l* and head *h*, and 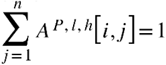. So, each amino acid at head *h* in layer *l* can attend to all other amino acids in the sequence, but the level of attention is determined by *A*^*P,l,h*^[*i,j*].

Based on the attention matrix 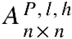, adjacency matrix 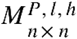 is constructed as follows:

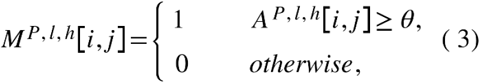

where the value of *θ* is determined by its application. We use attention and adjacency matrices for introducing our approaches to quantify representative heads in layers for some protein features (see Table 3).

#### 2.3.1 RLH_NNI algorithm to quantify the representative heads and layers of ProtAlbert for nearest-neighbor interaction

We propose the RLH_NNI algorithm to determine if head *h* in layer *l* of the ProtAlbert model represents the interaction of *k*-neighbor amino acids in dataset Δ. The main steps of this algorithm are defined as follows:

1. For each protein *P* ∈ Δ = {*P*_1_,…,*P*_*t*_},

i. The ***interaction of k-neighbor amino acids*** from the sequence *S*^*P*^ (│*S*^*P*^│=*n*) is quantified, as:

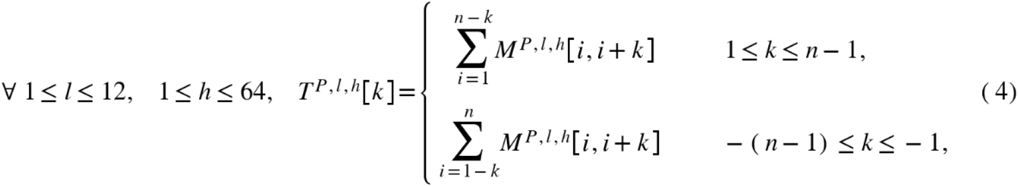

where adjacency matrix *M*^*P,l,h*^ is generated based on Eq.3 for each head *h* in layer *l*.
ii. The ***normalized k-neighbor interaction*** is defined like this:

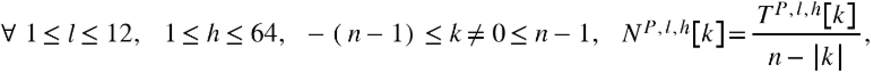

where │.│ shows the absolute function. For each head *h* in layer *l*, *N*^*P,l,h*^[*k*] indicates the percentage of positions in protein *P* which attend to the *k*^*th*^ amino acid in the neighbor.
iii. The ***weighted quantification of *k*-neighbor interaction*** is computed based on attention matrix, as:

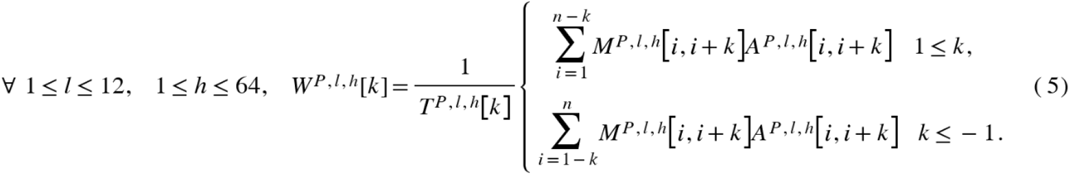
2. The ***average of normalized *k*-neighbor interaction*** is computed on dataset Δ, as:

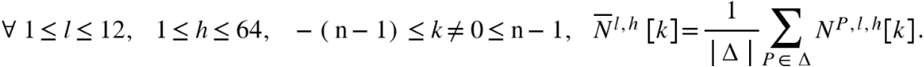
3. The maximum interaction value of neighbor amino acids at each head in the layer is computed to determine the ***nearest neighbor radius for interaction***, as:

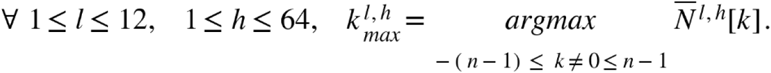
4. For each 1 ≤ *l* ≤ 12 and 1 ≤ *h* ≤ 64, if 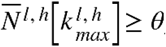,

i. Head *h* in layer *l* is announced ***representative for 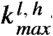-neighbor interaction*** on dataset Δ.
ii. For head *h* in layer *l*, the ***average of weighted quantification of 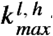-neighbor interaction*** is computed on dataset Δ, as:

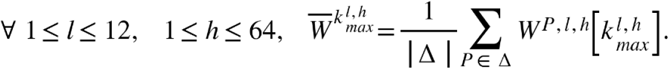

The average of weighted quantification 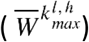 and the average of normalized interaction 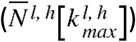 are compared to show the effect of discretizing of attention weights in the adjacency matrix.

#### 2.3.2 RH_SAA algorithm to quantify representative heads of ProtAlbert for specific amino acids

Here, we introduce the RH_SAA algorithm to investigate if the head *h* of ProtAlbert attends significantly to a specific amino acid in the protein dataset Δ. To quantify the quality of attention head *h* for amino acid *a* ∈ *AA*, we apply the ***F-measure criterion*** to evaluate the occurrence rate of amino acid versus the rest in this head. In the following, this algorithm is described in more detail:

1. For each protein *P* ∈ Δ = {*P*_1_,…, *P*_*t*_},

i. True positive, 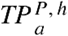, is defined based on the number of amino acid *a* in protein sequence 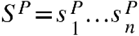 which is attended by at least one position of the sequence at head *h* in at least one layer:

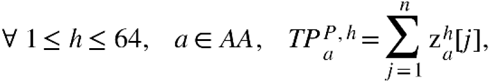

where

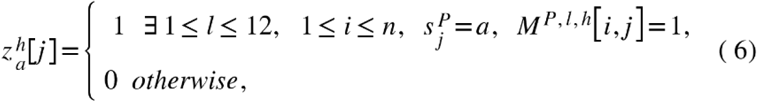

where adjacency matrix *M*^*P,l,h*^ is generated based on Eq.3 for head *h* in layer *l*.
ii. False positive, 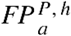, is obtained as follows:

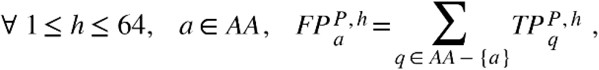

where 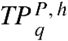 represents the number of amino acids *q* ≠ *a* attended by at least one sequence position at head *h* in at least one layer.
iii. False negative, 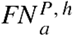, is computed as follows:

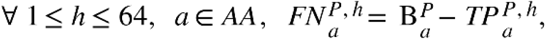

where 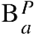 shows the frequency of amino acid *a* ∈ *AA* in sequence *S*^*P*^.
iv. ***F-measure criterion***, 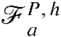, is computed to quantify head *h* for amino acid *a*:

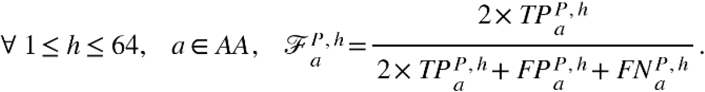
v. The ***relative occurrence of amino acid*** *a* of protein *P* in the head *h* is computed as:

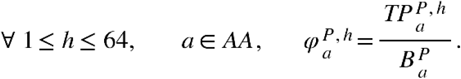
vi. The ***weighted occurrence of amino acid*** *a* of protein *P* is calculated as follows:

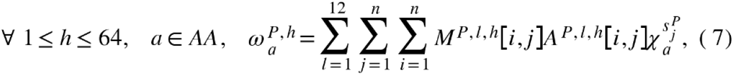

where

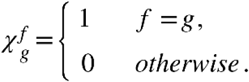
vii. The ***normalized weighted occurrence of amino acid*** *a* of protein *P* is calculated as follows:

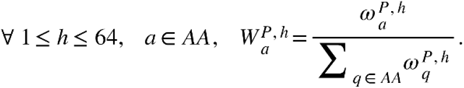
2. The ***average of F-measure*** is computed on the dataset Δ:

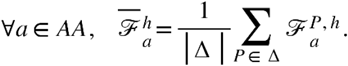
3. The ***candidate representative head*** for amino acid *a* is computed, as:

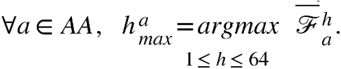
4. For each *a* ∈ *AA*, if 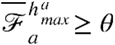:

i. Head 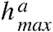 is announced as a ***representative head*** for amino acid *a*.
ii. In head 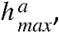, the ***average of normalized weighted occurrence of amino acid*** *a* is computed on dataset Δ, as:

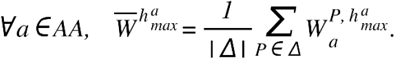

where the normalized weighted occurrence of amino acid *a* shows the effect of attention weights attending from each amino acid to *a* at head 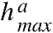.
iii. The ***average of the relative occurrence of amino acid*** *a* at head 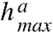 is computed on dataset Δ as:

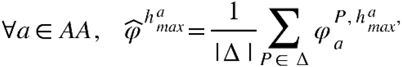

where the relative occurrence of amino acid *a* shows the probability of amino acid *a* detection at head 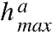.

#### 2.3.3 RH_BBP algorithm to quantify representative heads of ProtAlbert for biochemical and biophysical properties of amino acids

In this sub-section, we illustrate the RH_BBP algorithm to find representative heads of ProtAlbert on the biochemical and biophysical properties using the classification of amino acids based on the R-group. Table 1 shows this classification, 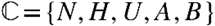. The algorithm is very similar to RH_SAA, which identifies specific heads for amino acids. In the following, the details of RH_BBP are available:

1. For each protein *P* ∈ Δ = {*P*_1_,…, *P*_*t*_},

i. In class 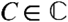, true positive is defined by 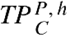 to show the number of amino acids from class *C* in protein sequence *S*^*P*^ (│ *S*^*P*^│ = *n*) attended by at least one position of the sequence at head *h* in at least one layer:

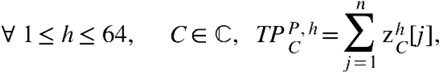

where

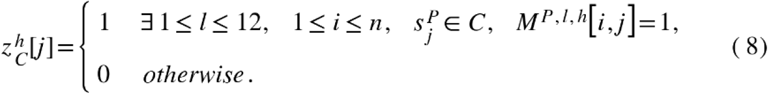
ii. For class 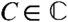, false positive in the head 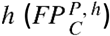 is obtained as follows:

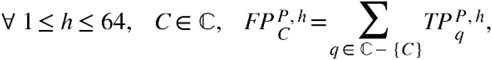

where 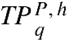 represents the number of amino acids from class *q* ≠ *C* attended by at least one sequence position *S*^*P*^ at head *h* in at least one layer.
iii. False negative, 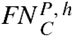, is computed as follows:

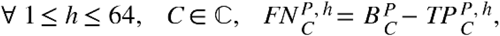

where 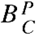 shows the frequency of the amino acids from class *C* in sequence *S*^*P*^.
iv. For class 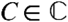, ***F-measure criterion***, 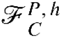, is computed to quantify head *h* at this class:

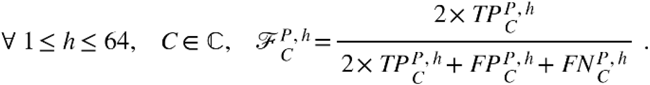
v. The ***relative occurrence of class*** *C* for protein *P* in head *h* is computed as:

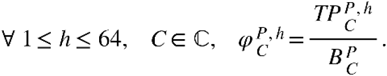
vi. The ***weighted occurrence of class*** *C* for protein *P* is calculated as follows:

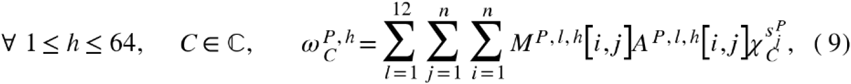

where

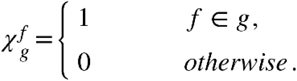
vii. The ***normalized weighted occurrence of class*** *C* for protein *P* is calculated as follows:

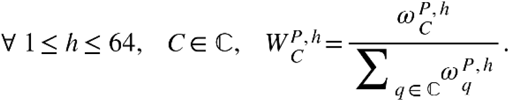
2. The ***average of F-measur***e is computed on dataset Δ:

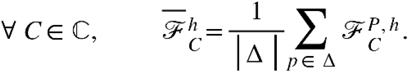
3. The ***candidate representative head*** for class *C* is calculated, as:

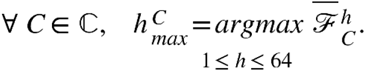
4. For each 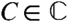, if 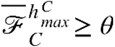:

i. Head 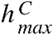 is announced as a ***representative head*** for class *C*.
ii. In head 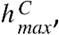, the ***average of normalized weighted occurrence of class*** *C* is computed on dataset Δ, as:

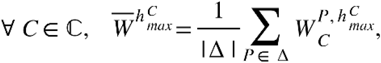

where the normalized weighted occurrence of class *C* shows the effect of attention weights attending from each amino acid to the amino acids in class *C* at head 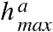.
iii. *The* ***average of the relative occurrence of class*** *C at head* 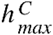 *on dataset* Δ *is computed as*:

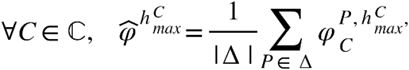

where the relative occurrence of class *C* shows the probability of class *C* detection at head 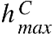.

#### 2.3.4 RH_PSS algorithm to quantify representative heads of ProtAlbert for protein secondary structure

Although ProtAlbert only has been pre-trained on protein sequences, we propose the RH_PSS algorithm on dataset Δ to assess attention heads about the protein secondary structure matrix (see Eq.2). The detail of this algorithm is as below:

1. For each protein *P* ∈ Δ = {*P*_1_,…, *P*_*t*_},

i. Predicting the secondary structure matrix 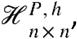, of protein *P* with length n for each head *h* as follows:

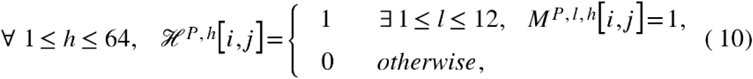

where adjacency matrix *M*^*P,l,h*^ is constructed based on Eq.3 for each head *h* in layer *l*.
ii. Making the natural secondary structure 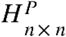 for protein *P* based on Eq.2.
iii. Computing the cosine similarity between 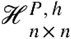 and 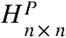 for each *h*, 1 ≤ *h* ≤ 64, cos(*ℋ*^*P,h*^, *H*^*P*^).
2. The average of cosine similarity is computed as:

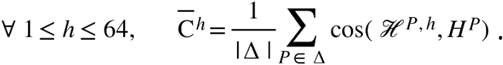

#### 2.3.5 RH_PTS algorithm to quantify representative heads of ProtAlbert for protein tertiary structure

We propose the RH_PTS algorithm to compare the natural protein contact map to the predicted contact map from head *h* on dataset Δ. The main steps of this algorithm are as follows:

1. For each protein *P* ∈ Δ = {*P*_1_,…, *P*_*t*_},

i. Making matrix 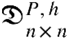 as:

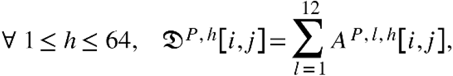

where *A*^*P,l,h*^ represents the attention matrix of protein *P* in layer *l* and head *h*.
ii. Normalizing matrix 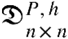 as bellow:

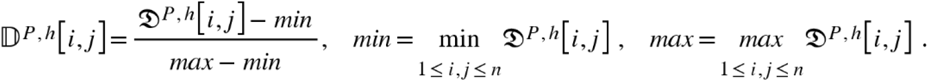
iii. Predicting contact map based on matrix 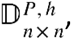, as:

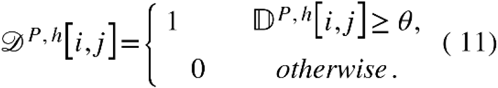
iv. Making real contact map 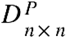 for protein *P* based on Eq.1.
v. Computing the cosine similarity between 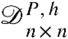 and 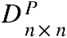 for each 1 ≤ *h* ≤ 64, 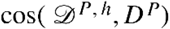.
2. Computing the average of cosine similarity as:

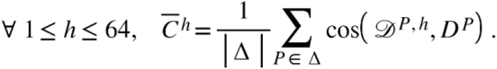
3. Finding head *h*_*max*_ to indicate the maximum similarity between natural and predicted contact maps:

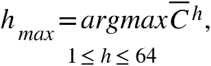

where head *h*_*max*_ is known as a representative head for contact maps.

### 2.4 Proposed algorithm for sequence profile prediction problem

In the second part of our work, we propose the PA_SPP algorithm for the sequence profile prediction problem. The input and output of this problem are defined as follows:

- **Input**: Sequence 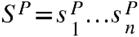 of protein *P*.
- **Output:** Predicting profile 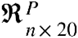 using pre-trained ProtAlbert.

To solve this problem, we apply pre-trained ProtAlbert to predict the masked token of an input sequence containing unknown amino acids in one position of the sequence *S*^*P*^. ProtAlbert model generates the most likely amino acids for that position. In other words, the model predicts the masked amino acid in the sequence based on the context of other amino acids surrounding it. This process is called masked token prediction and represented by:

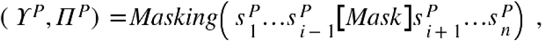

where generates two vectors 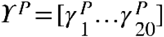 and 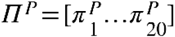. Vectors *ϒ*^*P*^ and *Π*^*P*^ represent the type of amino acids and the score for each amino acid replaced at the masked position in the sequence *S*^*P*^. For each 1 ≤ *j* ≤ 20, 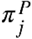 shows the score of substitution of amino acid 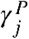 at position *i* of sequence *S*^*P*^. Figure 1 illustrates the PA_SPP algorithm for solving the profile prediction problem. In the first step, the sequence *S*^*P*^ is given as an input to the algorithm. In the second step, a zero-matrix named 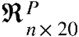 is defined where ℜ^*P*^[*i,j*] is updated during algorithm running by predicting the probability of *j*^*th*^ amino acid at the *i*^*th*^ position of sequence *S*^*P*^. The third step selects each position *i*, 1 ≤ *i* ≤ *n*, in sequence *S*^*P*^ for masking. In the fourth step, temporary memory *T* is defined to keep the sequence *S*^*P*^ with masking position *i*. In the fifth step, sequence *T* is fed to *Masking* process of ProtAlbert. The model generates two vectors ϒ^*P*^ and Π^*P*^ for position *i*. In the sixth step, we set the probability vector ϒ^*P*^ into the *i*^*th*^ row of matrix ℜ according to the order of amino acids in ϒ^*P*^. In the end, we call 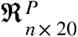 the predicted profile for protein *P*.

**Figure 1:**
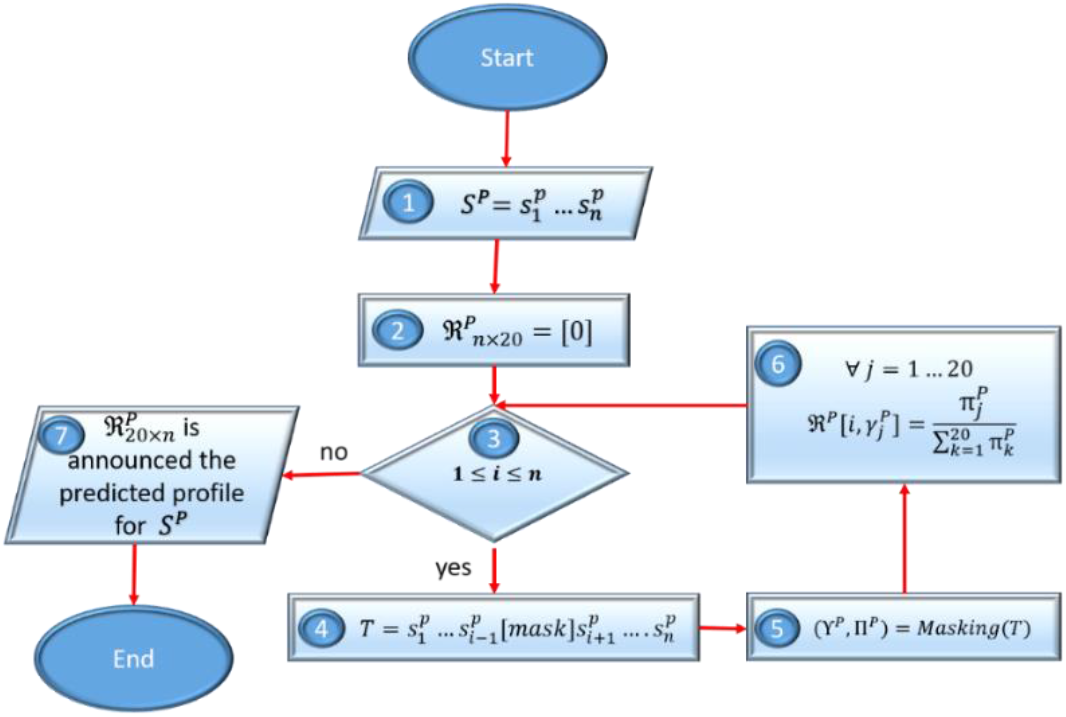
PA_SPP algorithm for protein profile prediction.

### 2.5 Dataset

In this study, we use the CASP13^x^ dataset. This dataset includes 194 proteins. We select 55 proteins (see Supplementary 1) whose profiles are available in the HSSP database. We call the selected proteins from CASP13, dataset Δ where | Δ | =55. The tertiary structure and sequence of each protein *P* ∈ Δ are extracted from the PDB database. In addition, their HSSP profiles are downloaded from xssp site.

The distribution of the extracted target sequences lengths is shown in Figure 2. In addition, Figure 3 represents the frequency of amino acids in the sequences of dataset Δ.

**Figure 2:**
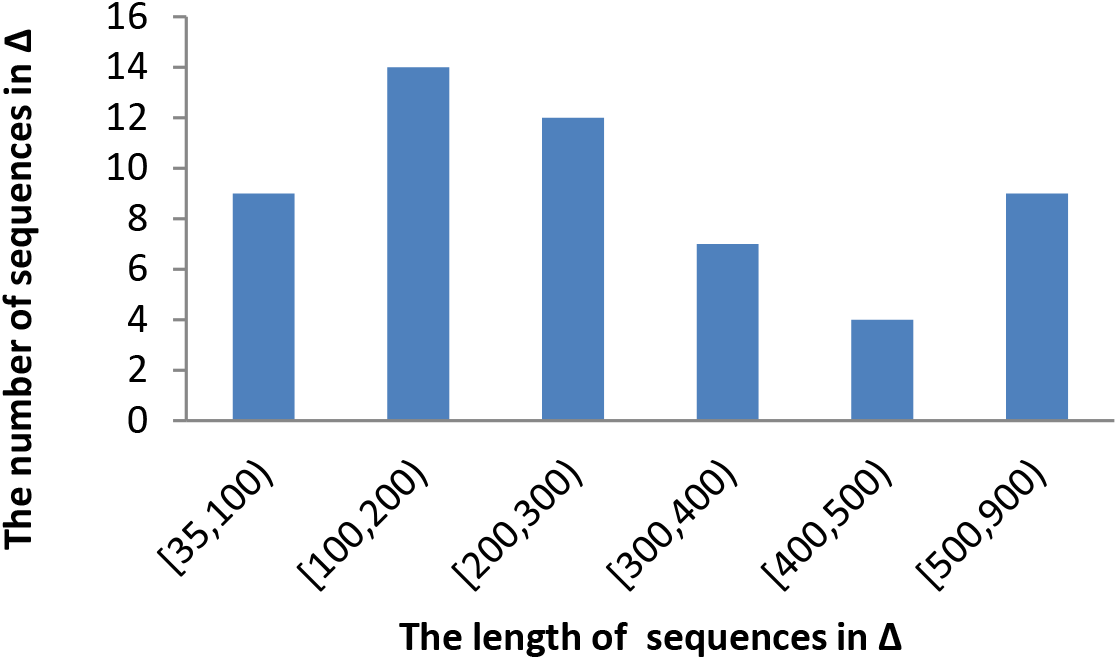
Distribution of the length of protein sequences in dataset Δ.

**Figure 3:**
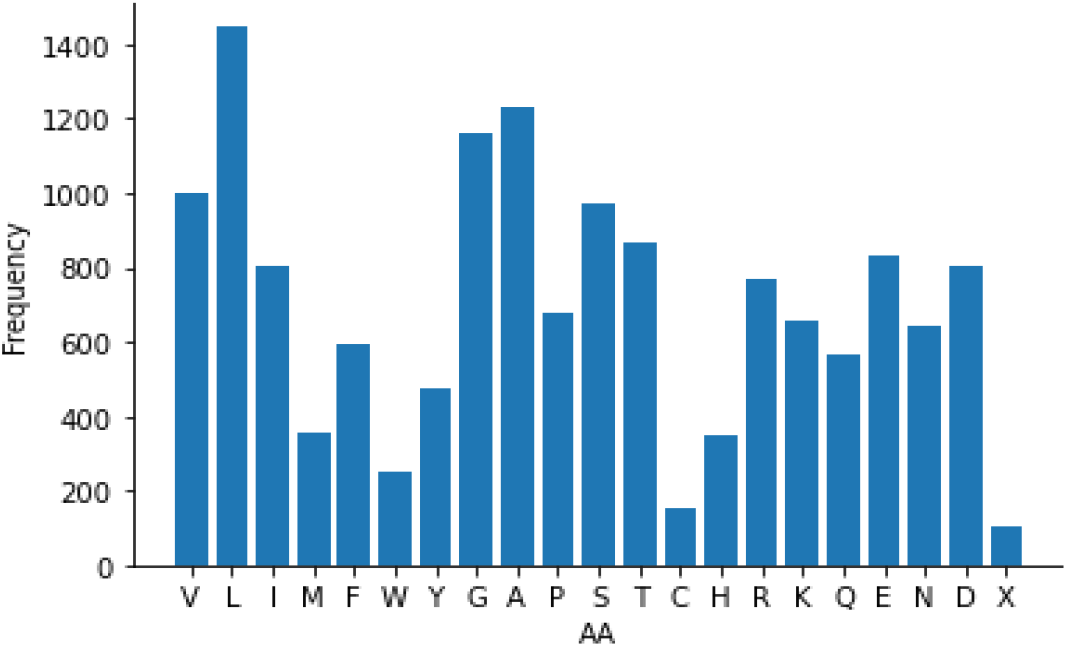
The frequency of amino acids (AA) in dataset Δ.

In addition to dataset Δ, we select three essential proteins (see Table 4) in different organisms for case studies to show that our result is generally reliable. The details of these proteins are available in Supplementary 2.

**Table 4:**
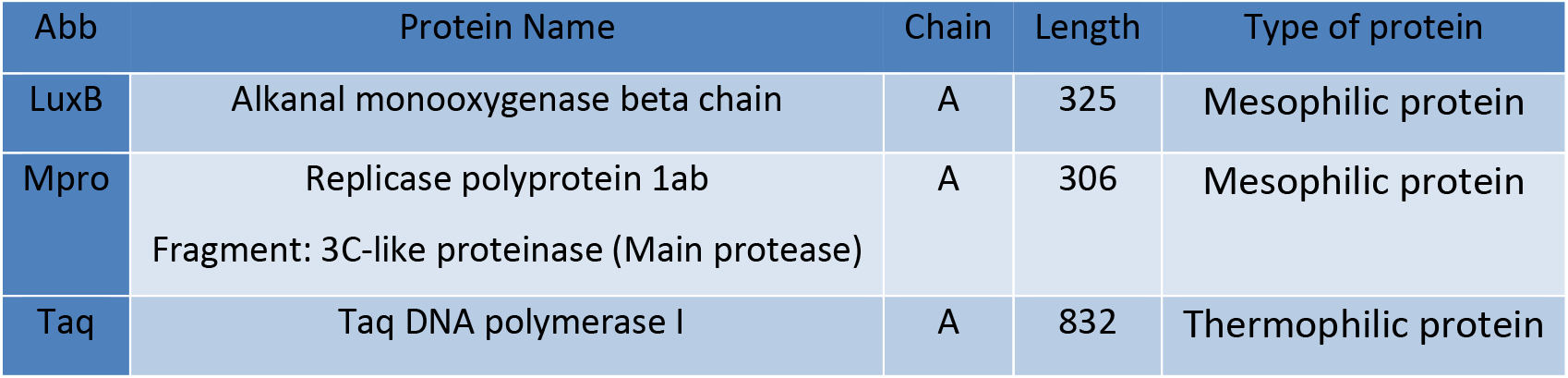
Details of three case study proteins.

## 3. Result and Discussion

In this section, we apply Δ ⊆ *CASP*13 and three case study proteins, LuxB, Mpro, and Taq, to analyze ProtAlbert as a pre-trained transformer on protein sequences. We find representative heads of ProtAlbert for five protein characteristics (Table 3). This part assures us that the heads contain the information required by a family of proteins. Then, we use this dataset for profile prediction. In the end, we compare the predicted profiles to the HSS profiles.

### 3.1 Analyzing ProtAlbert as a pre-trained transformer on protein sequences

Here, we find representative heads in the layers of ProtAlbert for five protein characteristics displayed in Table 3 using algorithms RLH_NNI, RH_SAA, RH_BBP, RH_PSS, and RH_PTS. In these algorithms, we use some cutoffs obtained by our trial and error. Cutoffs are set high for sequence feature analysis because ProtAlbert has been pre-trained on the protein sequences. For structures feature analysis, cutoffs are set low.

#### 3.1.1 Assessment of nearest-neighbor interactions at heads in layers of ProtAlbert

As mentioned in^42^, *k*-neighbor interaction where − 4 ≤ ≠ 0 ≤ 4 is known as proximal interaction, which is effective in the first step of protein folding. Here, we apply RLH_NNI algorithm on dataset Δ ⊆ CASP13 to find the representative heads in the layers of ProtAlbert for the nearest neighbor radius of amino acids interaction.

In Eq.4 and Eq.5 of this algorithm, we consider threshold 0.5 to make an adjacency matrix from the attention matrix. In the fourth step of RLH_NNI, we select representative head *h* in layer *l* for the interaction of 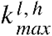-neighbor amino acids in dataset Δ, if 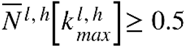. For each selected head *h* in layer *l* and protein *P* ∈ {*luxB*, *Mpro*, *Taq*}, 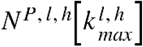 is computed. Table 5 shows the representative heads in layers for the interaction of 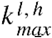-neighbor amino acids. The results show that the average of normalized 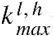-neighbor interaction 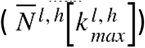 is close to the normalized 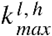-neighbor interaction on each protein 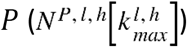 as case study.

**Table 5:**
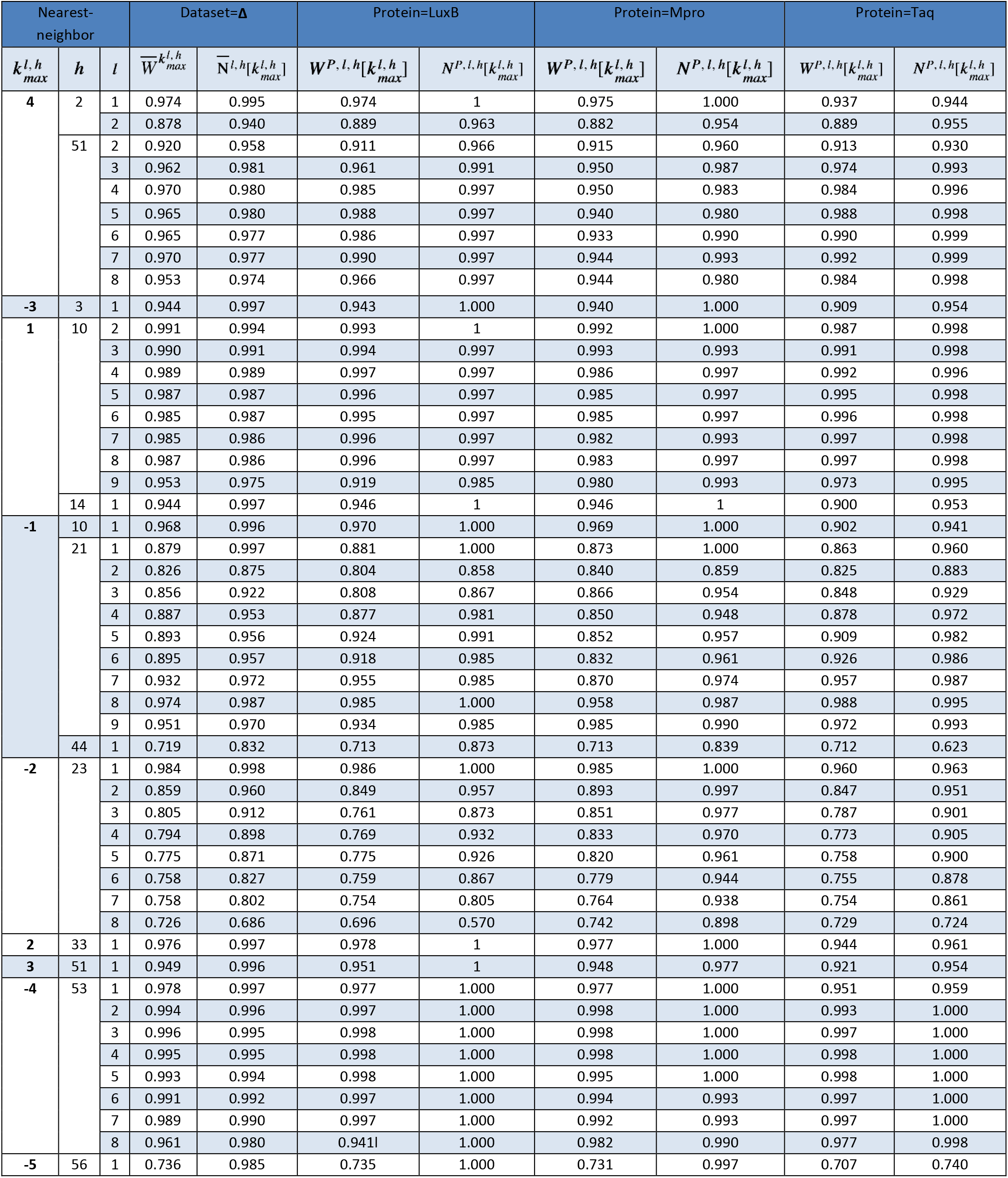
Representative heads in layers of ProtAlbert for nearest-neighbor interaction.

Also, the average of the weighted quantification of 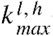-neighbor interaction, 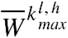 is calculated. Also, the weighted quantification for each case study protein *p*, 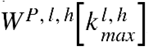, is available in this table. The values of *W* are close to *N* ones; it shows that attention weights are high in 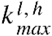-neighbor on the dataset and cases study proteins. The results show that

- head 10 in layers 2-9, head 21 in layers 1-9, and heads 14 and 44 in layer 1 represent interactions at one position apart.
- head 23 in layers 1-8 and head 33 in layer 1 are specific for interactions at two positions apart.
- heads 3 and 51 in layer 1 indicate interactions between each amino acid and its third neighbor in the sequence.
- head 51 in layers 2 – 8, head 53 in layers 1-8, head 2 in layers 1-2 represent the interaction between each amino acid and its fourth neighbor in the sequence.
- head 56 in layer 1 is specific for interactions at five positions apart.

In conclusion, we have identified the representative heads in different layers for proximal positions in proteins. According to ^42^, proximal positions are essential in the first step of protein folding.

#### 3.1.2 Assessment of the type of amino acids at heads of ProtAlbert

In this sub-section, we use the RH_SAA algorithm to find a representative head for each amino acid on dataset Δ ⊆ CASP13. In Eq.6 and Eq.7 of this algorithm, we consider threshold 0.4 to make adjacency matrix from attention matrix.

In the third step of RH_SAA, we select candidate representative head 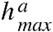 for amino acid *a*. At the fourth step, head 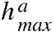 is announced as a representative head for amino acid *a*, if 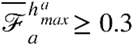. Meanwhile, we compute the F-measure criterion, 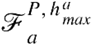, for amino acid *a* in each protein *P* ∈ {*LuxB*, *Mpro*, *Taq*} at head 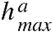. Table 6 shows that the average of the F-measure is similar to the F-measure of each case study.

**Table 6:** The representative heads for amino acids found based on F-measure 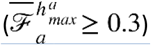.

| Head | Amino acid | Dataset= $\Delta$ | | | Protein=LuxB | | | Protein=Mpro | | | Protein=Taq | | |
| --- | --- | --- | --- | --- | --- | --- | --- | --- | --- | --- | --- | --- | --- |
| | | $\overline{\mathcal{F}}_a^{h_{max}^a}$ | $\widehat{\varphi}_{max}^{h_{max}^a}$ | $\overline{W}_{max}^{h_{max}^a}$ | $\mathcal{F}_a^{P, h_{max}^a}$ | $\varphi_a^{P, h_{max}^a}$ | $W_a^{P, h_{max}^a}$ | $\mathcal{F}_a^{P, h_{max}^a}$ | $\varphi_a^{P, h_{max}^a}$ | $W_a^{P, h_{max}^a}$ | $\mathcal{F}_a^{P, h_{max}^a}$ | $\varphi_a^{P, h_{max}^a}$ | $W_a^{P, h_{max}^a}$ |
| 8 | E,<br>D | 0.41,<br>0.41 | 0.85,<br>0.89 | 0.34,<br>0.40 | 0.45,<br>0.48 | 0.90,<br>0.90 | 0.42,<br>0.46 | 0.20,<br>0.44 | 0.78,<br>1.0 | 0.24,<br>0.49 | 0.59,<br>0.39 | 0.78,<br>0.85 | 0.53,<br>0.33 |
| 13 | P | 0.52 | 0.9 | 0.39 | 0.48 | 1 | 0.32 | 0.61 | 1.0 | 0.41 | 0.47 | 0.72 | 0.29 |
| 18 | D | 0.42 | 0.97 | 0.56 | 0.48 | 1 | 0.6 | 0.43 | 1.0 | 0.56 | 0.46 | 0.9 | 0.6 |
| 20 | W | 0.57 | 0.94 | 0.58 | 0.44 | 1 | 0.67 | 0.75 | 1.0 | 0.57 | 0.81 | 0.92 | 0.86 |
| 63 | H | 0.32 | 0.5 | 0.37 | 0.31 | 0.3 | 0.42 | 0.44 | 0.57 | 0.48 | 0.37 | 0.44 | 0.38 |

Moreover, this table represents the average relative occurrence of amino acid *a* in dataset Δ and each case study protein *P* ∈ {*LuxB*, *Mpro*, *Taq*} by 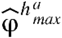 and 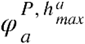, respectively. In addition, the average of normalized weighted occurrence of amino acid *a* in dataset Δ and case study protein *P* are shown by 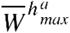 and 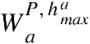, respectively. As a result, we find that

- the average of F-measure on the dataset is close to case study ones
- heads 8 and 18 can support hydrophilic acidic amino acids, aspartic acid (D) and glutamic acid (E)
- heads 13, 20, and 63 are specific for proline (P), tryptophan (W), and histidine (H), respectively.

To better understand the selected heads for specific amino acids, Figure 4 shows the weighted stacking of amino acids 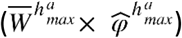 at attention heads 8, 13, 18, 20, and 63.

**Figure 4:**
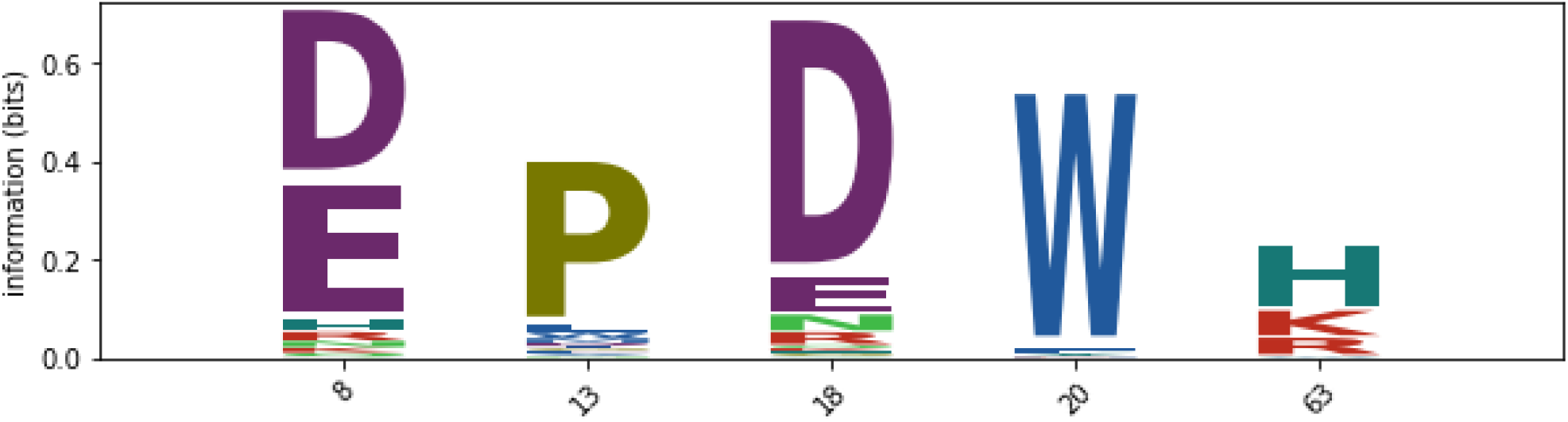
Logo consists of the weighted stacking of amino acids relative to the occurrences of amino acids in the protein sequences at heads 8, 13, 18, 20, and 63.

#### 3.1.3 Assessment of biochemical and biophysical properties of amino acids at heads of ProtAlbert

In the previous sub-section, we found representative heads 8 and 18 for amino acids D and E. They are hydrophilic acidic amino acids. In the following, we assess the heads in layers to find more biochemical and biophysical properties based on the R-group of amino acids. This classification, 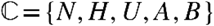, is shown in Table 1. To do the assessment, we apply the RH_BBP algorithm on dataset Δ ⊆ CASP13. In Eq.8 and Eq.9 of this algorithm, we consider threshold 0.4 to make adjacency matrix from attention matrix. In the third step of RH_BBP, head 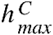 is selected to identify the maximum quantity for class 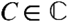. At the fourth step, we announce that head 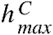 is representative for class *C* if 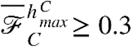. Meanwhile, we compute 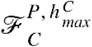 for each protein *P* ∈ {*luxB*, *Mpro*, *Taq*} at the selected head 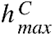. Table 7 shows that the average F-measure for class *C* at representative head 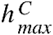 is similar to the case study ones.

**Table 7:** The representative heads for the classes in set ℂ based on F-measure 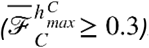.

| Head | Class Amino acids | Dataset= $\Delta$ | | | Protein=LuxB | | | Protein=Mpro | | | Protein=Taq | | |
| --- | --- | --- | --- | --- | --- | --- | --- | --- | --- | --- | --- | --- | --- |
| | | $\overline{\mathcal{F}}_C^{h_{max}^C}$ | $\widehat{\varphi}^{h_{max}^C}$ | $\overline{W}^{h_{max}^C}$ | $\mathcal{F}_C^{P, h_{max}^C}$ | $\varphi_C^{P, h_{max}^C}$ | $W_C^{P, h_{max}^C}$ | $\mathcal{F}_C^{P, h_{max}^C}$ | $\varphi_C^{P, h_{max}^C}$ | $W_C^{P, h_{max}^C}$ | $\mathcal{F}_C^{P, h_{max}^C}$ | $\varphi_C^{P, h_{max}^C}$ | $W_C^{P, h_{max}^C}$ |
| 8 | A={E,D} | 0.66 | 0.86 | 0.74 | 0.72 | 0.81 | 0.88 | 0.53 | 0.92 | 0.61 | 0.77 | 0.81 | 0.86 |
| 43 | B={R,H,K} | 0.46 | 0.64 | 0.28 | 0.51 | 0.71 | 0.33 | 0.36 | 0.69 | 0.15 | 0.56 | 0.60 | 0.33 |
|  | U={S,T,C,N,Q} | 0.40 | 0.40 | 0.44 | 0.46 | 0.45 | 0.43 | 0.58 | 0.56 | 0.60 | 0.14 | 0.37 | 0.35 |
| 44 | N={G,A,V,L,I,P,M} | 0.63 | 0.99 | 0.67 | 0.56 | 1 | 0.58 | 0.62 | 1 | 0.59 | 0.71 | 0.97 | 0.73 |
| 49 | H={F,Y,W} | 0.45 | 0.86 | 0.5 | 0.5 | 0.83 | 0.46 | 0.54 | 0.94 | 0.48 | 0.46 | 0.86 | 0.33 |

Moreover, this table represents the average relative occurrence of class *C* for dataset Δ and each case study protein *P* by 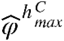 and 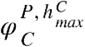, respectively. In addition, the average weighted occurrence of class *C* and each case study protein *P* at this head is shown by 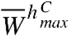 and 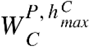, respectively.

In conclusion, representative heads 8, 44, and 49 show hydrophilic acidic, hydrophobic aliphatic, and hydrophobic aromatic amino acids, respectively. Also, head 43 can represent both polar and basic amino acids. Figure 5 consists of the weighted stacking of amino acids 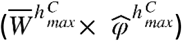 at attention heads 8, 43, 44, and 49.

**Figure 5:**
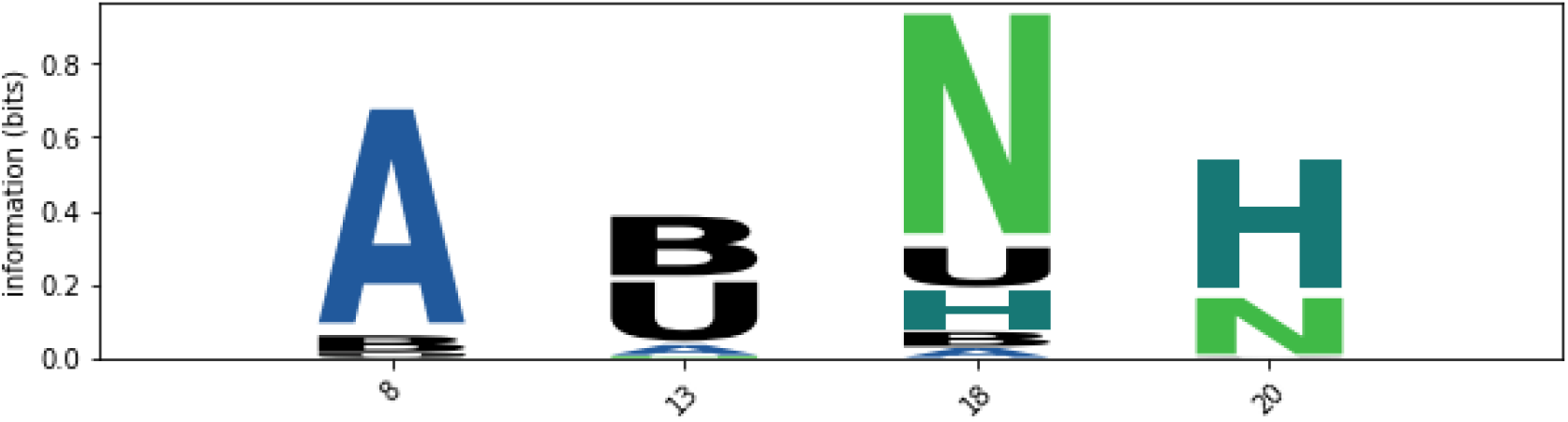
The logo consists of weighted stacking of amino acids in class 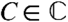 relative to the occurrences of these amino acids in the protein sequences at heads 8, 43, 44, and 49.

#### 3.1.4 Assessment of the protein secondary structure at heads of ProtAlbert

ProAlbert has been pre-trained on protein sequences, but we use the RH_PSS algorithm on dataset Δ ⊆ CASP13 to show that some heads with high attention weights are attending from helix to helix, sheet to sheet, and coil to coil. In Eq.10 of this algorithm, we consider threshold 0.1 to make an adjacency matrix from the attention matrix. At step two of RH_PSS, the average of cosine similarity, 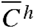, between the predicted and natural secondary structure matrices is computed.

Figure 6 shows the heatmaps of 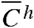, at each head *h*, 1 ≤ *h* ≤ 64 on data set Δ ⊆ *CASP*13. In addition, the cosine similarity, cos(*ℋ*^*P,h*^, *H*^*P,h*^), for each case study protein *P* ∈ {Taq, Mpro, LuxB}is computed. The high similarity between the predicted and natural protein secondary matrices can be seen at heads 2, 3, 8, 9, 10, 13,14, 18, 20, 21, 23, 32, 49, 51, 53, 56, and 63. Some of these heads are common with the heads in nearest-neighbor interaction. After removing the common heads, we find that heads 8, 9, 13,18, 20, 32, 49, and 63 are only informative about the secondary structure. These heads show more attention from the secondary structure of each amino acid to the same structure, with less than 6 amino acids in neighbors.

**Figure 6:**
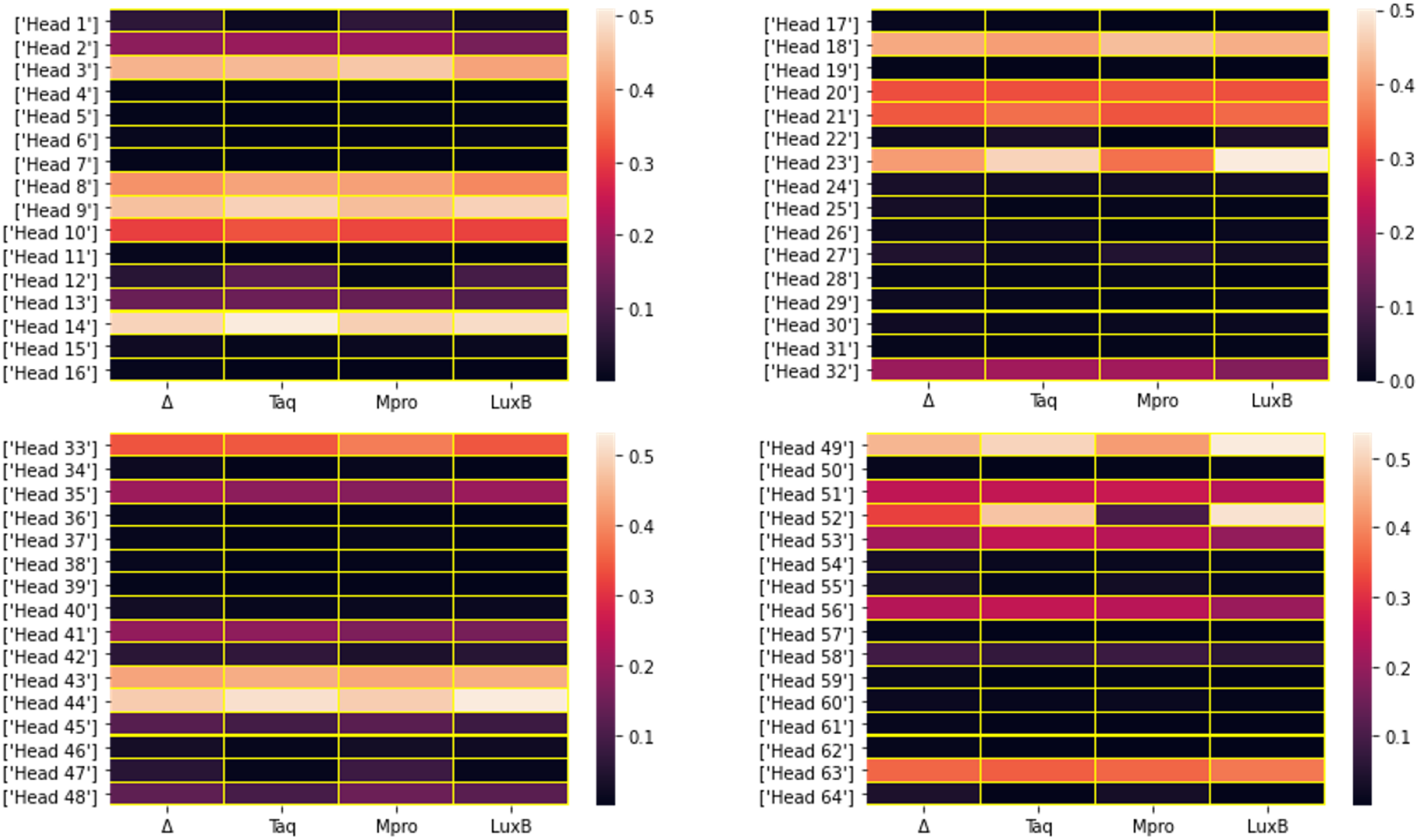
Heatmap of cosine similarity between the predicted and natural protein secondary structure matrices

#### 3.1.5 Assessment of the protein tertiary structure at heads of ProtAlbert

In this sub-section, we compute the similarity between the natural contact map (see Eq.1) and predicted contact map of protein *P* using the RH_PTS algorithm on dataset Δ ⊆ CASP13. In this algorithm, threshed 0.1 is defined for Eq.11 to discretize the predicted contact map.

Table 8 shows the average similarity between the predicted and natural contact maps at *h*_*max*_ = 10 on dataset Δ obtained from step three of the algorithm. Then 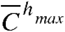 is calculated based on the second step of RH_PTS. In addition, the cosine similarity, 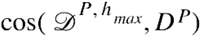 is computed for each case study protein *P* ∈ {*Taq, Mpro, LuxB*}. It seems that head 10 can show appropriate information on contact maps.

**Table 8:** Cosine similarity between natural and predicted contact map for proteins at head 10.

| Data | Cosine similarity |
| --- | --- |
| Dataset=$\Delta$ | $\bar{C}^{10}_{max}=0.7949$ |
| P=Mpro | $\cos(\mathcal{D}^{P, 10}, D^P)=0.7299$ |
| P=Taq | $\cos(\mathcal{D}^{P, 10}, D^P)=0.8071$ |
| P=LuxB | $\cos(\mathcal{D}^{P, 10}, D^P)=0.8167$ |

### 3.2 Predicting profile using ProtAlbert

The above assessment shows that transformers can capture some protein features from a single sequence to represent the protein family. These features can lead us to find appropriate information about the homologous sequences of each single protein sequence given as an input to ProtAlbert. Therefore, the PA_SPP algorithm (see Figure 1) employs pre-trained ProtAlbert to predict a profile for a query sequence. Here, we compare the predicted profiles to real ones obtained from the homologous sequences (HSSP profile).

For each protein *P* ∈ ⊆ CASP13 and three case study proteins, Taq, Mpro, and LuxB, the PA_SPP algorithm predicts profile ℜ^*P*^. Then, we compare the similarity of the predicted profile ℜ^*P*^ to the HSSP profile *R*^*P*^ using cosine similarity. We want to show that the predicted profile is close to the HSSP profile. It should be noted; some HSSP profiles are more reliable than the other ones because the number distribution of sequences aligned to the query sequence is different. For example, some profiles are obtained by less than 100 aligned sequences, and some are made based on more than 1000 aligned sequences. Therefore, the HSSP profile constructed with more aligned sequences is more reliable. So, the weighted average similarity between predicted and HSSP profiles are computed by the number of aligned sequences. Figure 7 shows that the predicted profiles are more similar to the HSSP profiles with more alignment sequences.

**Figure 7:**
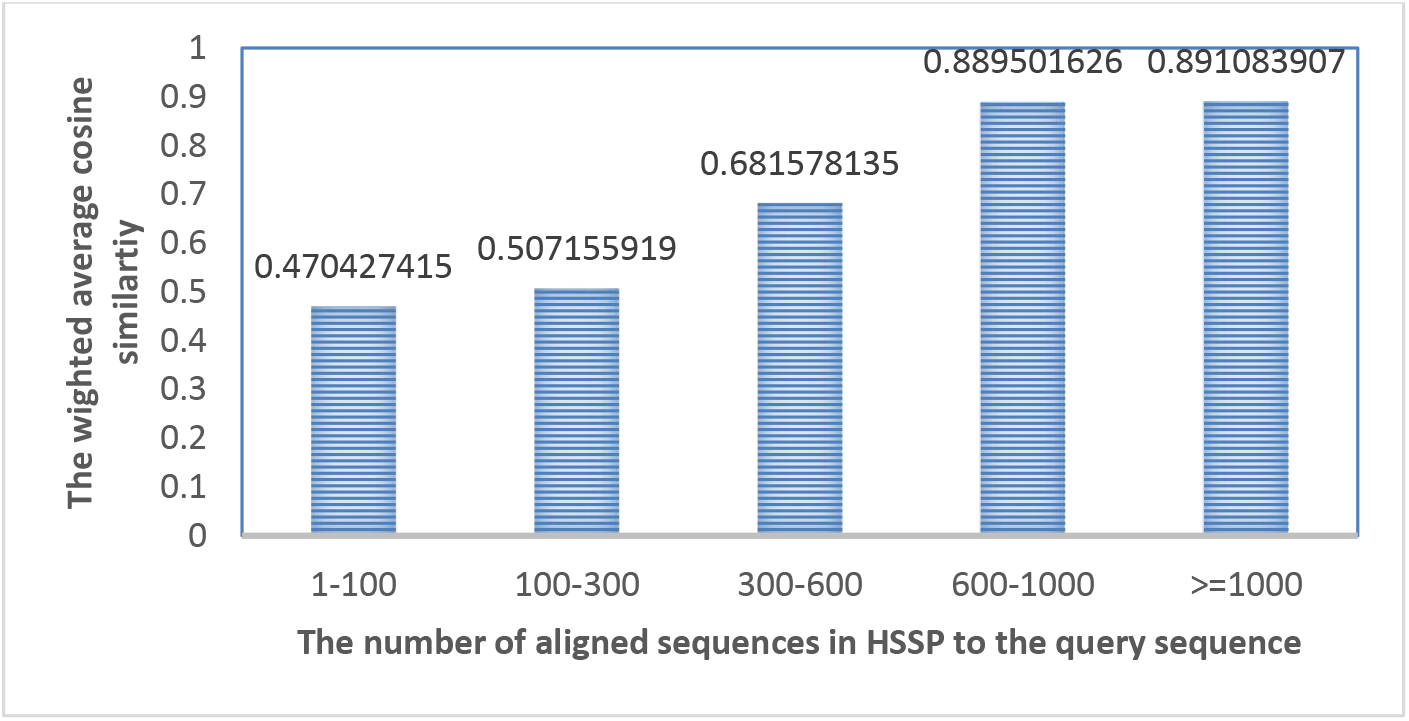
The weighted cosine similarity between predicted profiles and HSSP profiles based on the number of aligned sequences to the query sequence.

## 4. Conclusion

This paper contained two parts, ProtAlbert model analysis and profile prediction. Most previous studies used pre-trained transformer models to generate an embedding for protein sequence in different bioinformatics problems as a black box. Here, we would like to find the representative heads in layers for some protein characteristics. For this assessment, we used ProtAlbert because its efficiency enables us to run the model on longer sequences with less computation power while having similar performance with the other pre-trained transformers on proteins which is a great advantage for us.

In this study, we did not train or fine-tune ProtAlbert. In other words, we used pre-trained ProtAlbert to determine the interaction of nearest-neighbor amino acids, type of amino acids, biochemical and biophysical properties of amino acids, protein secondary structures, and tertiary structures at attention heads in different layers. This analysis is crucial because it shows that ProtAlbert captures some protein family features from only sequences. It led us to propose an algorithm called PA_SPP for profile prediction from a query sequence using ProtAlbert. The results showed that the predicted profile is close to the profile obtained from the homologous sequences.

We believe that the proposed algorithm for profile prediction can help the researchers to make a profile for a query sequence while there are no similar sequences to the query sequence in the database. In the future, we can improve this predictor with new transformer models.

## Footnotes

i **R**epresentative **L**ayers and **H**eads of ProtAlbert for **N**earest **N**eighbor **I**nteractions

ii **R**epresentative **H**eads of ProtAlbert for **S**pecific **A**mino **A**cids

iii **R**epresentative **H**ead of ProtAlbert for **B**iochemical and **B**iophysical **P**roperties of amino acids

iv **R**epresentative **H**eads of ProtAlbert for **P**rotein **S**econdary **S**tructure

v **R**epresentative **H**eads of of ProtAlbert for **P**rotein **T**ertiary **S**tructure

vi Using **P**rot**A**lbert for **S**equence **P**rofile **P**rediction

vii https://microbenotes.com/amino-acids-properties-structure-classification-and-functions/

viii https://www.rcsb.org/

ix https://www3.cmbi.umcn.nl/xssp/

x https://www.predictioncenter.org/casp13/domains_summary.cgi

